# High resolution live imaging of tardigrade response to anoxia

**DOI:** 10.1101/2025.01.22.634244

**Authors:** Anna-Mari Haapanen-Saaristo, Sara Calhim, Ilkka Paatero

## Abstract

Tardigrades are well-known for their ability to tolerate extreme environmental conditions such as heat, drought and lack of oxygen by undergoing cryptobiosis. The molecular responses to stress have been studied in detail, but the physiological and morphogenetic changes during cryptobiosis are less understood. We developed new live high-resolution fluorescence microscopy protocols to visualize the tardigrade response to lack of oxygen – anoxybiosis. High-resolution time-lapse imaging enabled analysis of cellular morphology and tracking of cell movements during anoxybiosis. These analyses revealed considerable changes in morphology, composition and movement of storage cells. Our observations and new imaging protocols can be used to study morphological and cellular response to stress in tardigrades.

## Introduction

Tardigrades are small aquatic animals, which are known for their ability to withstand extreme conditions^1–3^. It is notable that these organisms demonstrate remarkable resilience to stress when compared to other animal groups. In contrast to larger organisms, which possess the capacity for extensive migration, small organisms, including tardigrades, are constrained by limited mobility and are therefore compelled to adapt and endure environmental changes. These unique mechanisms that the tardigrades use for survival are under on-going investigation.

The ability of tardigrades to survive extreme environmental stressors is based on cryptobiotic states, where metabolic activity is reduced to near-undetectable levels^4^. The phenomenon of cryptobiosis was first documented already in the late 1950s ^5^. One of these states is anoxybiosis, triggered by a lack of oxygen, which differs from other well-known cryptobiotic forms, such as anhydrobiosis, where survival is facilitated by desiccation. Unlike anhydrobiosis, where tardigrades can remain in a dry, dormant state for years, anoxybiosis is a temporary state of suspended animation that occurs under conditions of severe oxygen deprivation^6^. Once favorable conditions are restored, tardigrades can rapidly recover, resuming normal metabolism and activity^7^.

Investigating anoxybiosis in tardigrades offers unique insights into the cellular and molecular adaptations that allow organisms to endure hypoxic or anoxic conditions. Compared to other cryptobiotic forms, the mechanisms underlying anoxybiosis are less well understood, and while some studies have focused on tardigrades’ ability to survive desiccation, far fewer have explored their response to prolonged oxygen deprivation^6,7^. Responses to anoxic conditions vary among different organisms, including invertebrates such as several species of nematodes^8^. During oxygen deprivation, some other invertebrates may undergo changes in cellular morphology, including alterations in cellular shape, size, and structure^9^. These changes are often linked to physiological adaptations, such as reduced metabolic rates and changes in ion transport^9–11^.

Understanding these dynamics could illuminate broader ecological implications, particularly as climate change continues to alter habitats, potentially exposing tardigrades to unprecedented combinations of environmental challenges. The elucidation of stress responses of tardigrades has been also proposed to provide novel views of developing medical treatments and therapies^3,12,13^.

In this study, our aim was to develop high-resolution fluorescence microscopy methodology for a detailed analysis of the tardigrade’s transition into anoxybiosis.

## Results

### Protocol for live imaging of tardigrade anoxybiosis

Anoxybiosis is species specific adaptation mechanism and there are tardigrade species that tolerate anoxia poorly^6,14^. The range of toleration can vary from hours to several days^14^. To find a suitable model for anoxybiosis studies we tested two eutardigrade species *Macrobiotus ripperi* (Stec, Vecchi & Michalczyk, sp. nov. 2021) and *Paramacrobiotus fairbanksi (*Schill, Förster, Dandekar & Wolf, 2010) (Fig. 1A). *P. fairbanksi* is a larger species and *M. ripperi* is smaller (Fig. 1B). *M. ripperi* was able to withstand anoxia better than *P. fairbanksi* (Fig. 1C). In addition to its better tolerance of low oxygen levels, *M. ripperi* was more transparent which was beneficial for the imaging experiments. Some tardigrade species have significant cuticular autofluorescence, which can interfere with fluorescence imaging of internal structures^15^. *P. fairbanksi* had stronger autofluorescence than *M. ripperi* in cuticular structures and was opaquer, which was observed as whiter shade (Fig. 1A). These observations were consistent with earlier findings in *Paramacrobiotus sp*.^16^. Therefore, we decided to focus our study on just *M. ripperi*.

**Figure 1.**
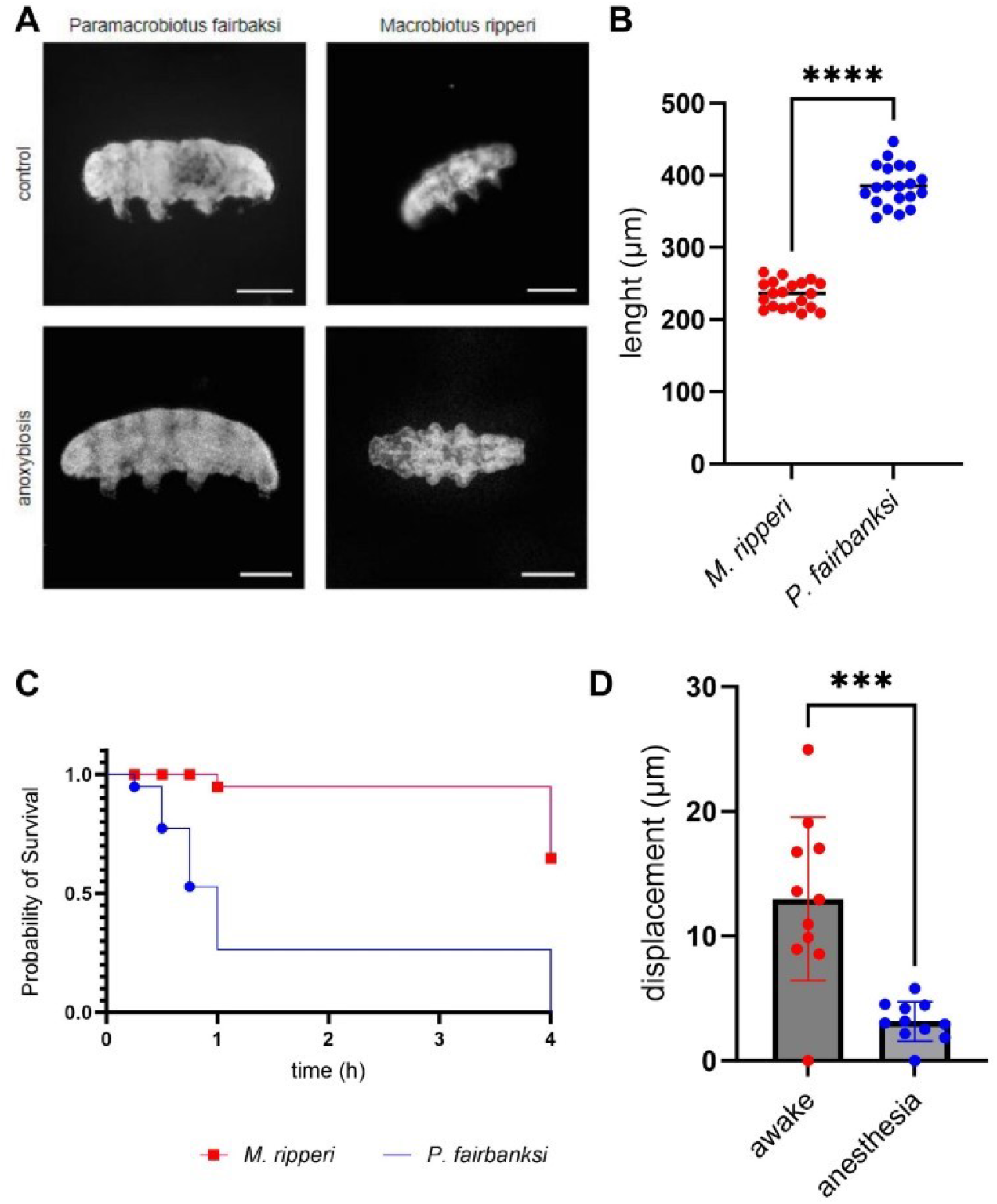
Finding suitable conditions for imaging studies of anoxia response in tardigrades. (A) Reflected light stereomicroscopy images of tardigrades. Anoxybiosis affected the size of tardigrade. Scale bar 100 μm. (B) Measurement of size of tardigrades. The two species (M.ripperi and P.fairbanksi) were of different size (n_ripperi = 20, n_fairbanksi=20, Mann-Whitney P<0.0001). (C) Analysis of survival under anoxia. M. ripperi showed better survival rates compared to P. fairbanksi. P.fairbanksi did not correspond to chemical hypoxia well, leading to 100% dead rate after 4-hour exposure. n_ripperi = 19, n_fairbanksi = 19. Log-rank (Mantel-Cox) test P <0.001. (D) Analysis of tardigrade movement during live imaging. Short bright-field time-lapse movies or awake and anaesthetized tardigrades were obtained with a stereomicroscope. There was significant decrease in motility when using Tricaine (MS222) as anesthetic. Mann-Whitney unpaired t-test P<0.0004, n_ripperi = 5, n_fairbanksi =5, with 11 measurement points each.

To enable live imaging studies, we needed to develop methods for immobilization of live tardigrades. Embedding of samples into low-melting point agarose is widely used to immobilize small organisms for imaging^17,18^. When we embedded *M. ripperi* in agarose, we observed that agarose alone could not restrain movement of *M. ripperi* sufficiently. Therefore, we tested MS-222 used in zebrafish embryo anesthesia^19^ and levamisole, which is used in immobilization of *C. elegans* for live imaging^20^. Levamisole proved lethal to *M. ripperi*, but MS-222 was well tolerated and significantly reduced the movement of *M. ripperi* (Fig. 1, supplemental movies S1 and S2). The MS-222 anesthesia together with mounting in low-melting point agarose provided sufficient control for high-resolution imaging experiments.

### Fluorescence staining of live tardigrades

In order to analyze the morphological changes of tardigrades during anoxybiosis, we aimed to identify suitable fluorescent labels. Although the use of genetically encoded fluorescent reporters has been recently demonstrated in some other tardigrade species^21^, we explored organic vital fluorescent dyes due to easier implementation and flexibility in experimentation. Two dyes previously employed for vital staining of other organisms, acridine orange and Nile red, were observed to label discrete structures in *M. ripperi* (Fig. 2A) with sufficient intensity. The intensity was significantly increased upon long incubations (Fig. 2B), most likely due to slow absorption of dye through chitin-rich outer cuticle of tardigrades. Acridine orange stains nucleic acids and Nile red stains lipids^22,23^.

**Figure 2.**
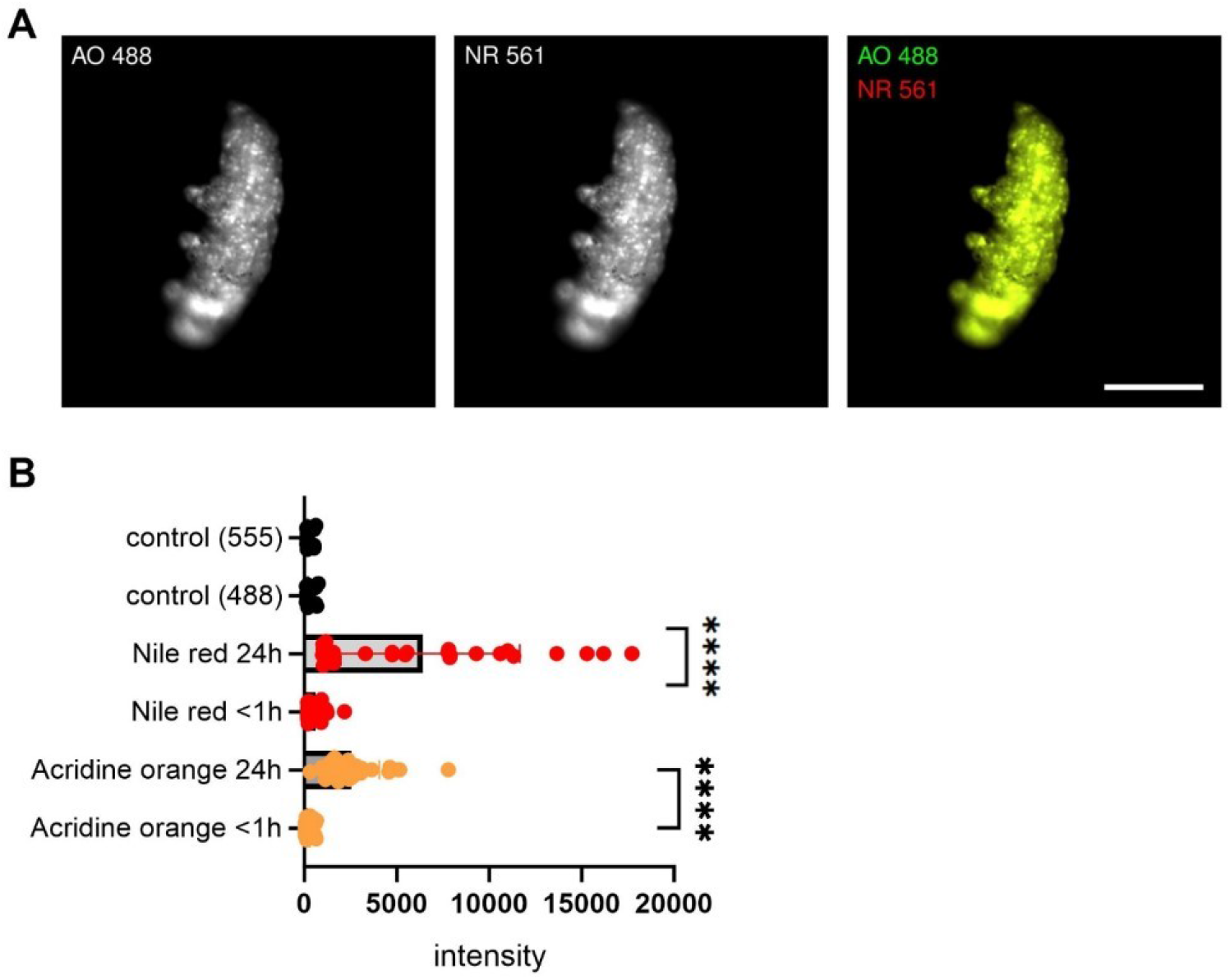
Vital fluorescent labeling of *M. ripperi*. **(A)** Fluorescent microscopy images of *M. ripperi* stained with acridine orange (AO) and Nile Red (NR) imaged with AxioZoom stereomicroscope. Scale bar 100 μm. **(B)** Quantification of fluorescence intensity. Background is reduced from actual fluorescent measurements (control). Number of tested animals: n(control) = 38, n(NR24h) = 25, n(NR1h) = 27, n(AO24h) = 23, n(AO1h) = 33. Unpaired t-test, AO P<0.0001 and NR P<0.0001.

### Tardigrades exhibit rapid changes within the body upon anoxia

The morphology of tardigrades is altered in anoxybiosis^14^. To analyze temporal changes in response to low oxygen levels, we chemically depleted oxygen from the culture medium and imaged *M. ripperi* during transition in anoxybiosis (Fig. 3A). The body length was increased during anoxia and visible increase was observed already in the early stages of anoxic exposure. The fastest rate of body length change was after 25 minutes (Fig. 3B). The volumetric measurements indicated that the alteration was not merely an increase in length, but rather a generalized expansion of the body (Fig. 3C).

**Figure 3.**
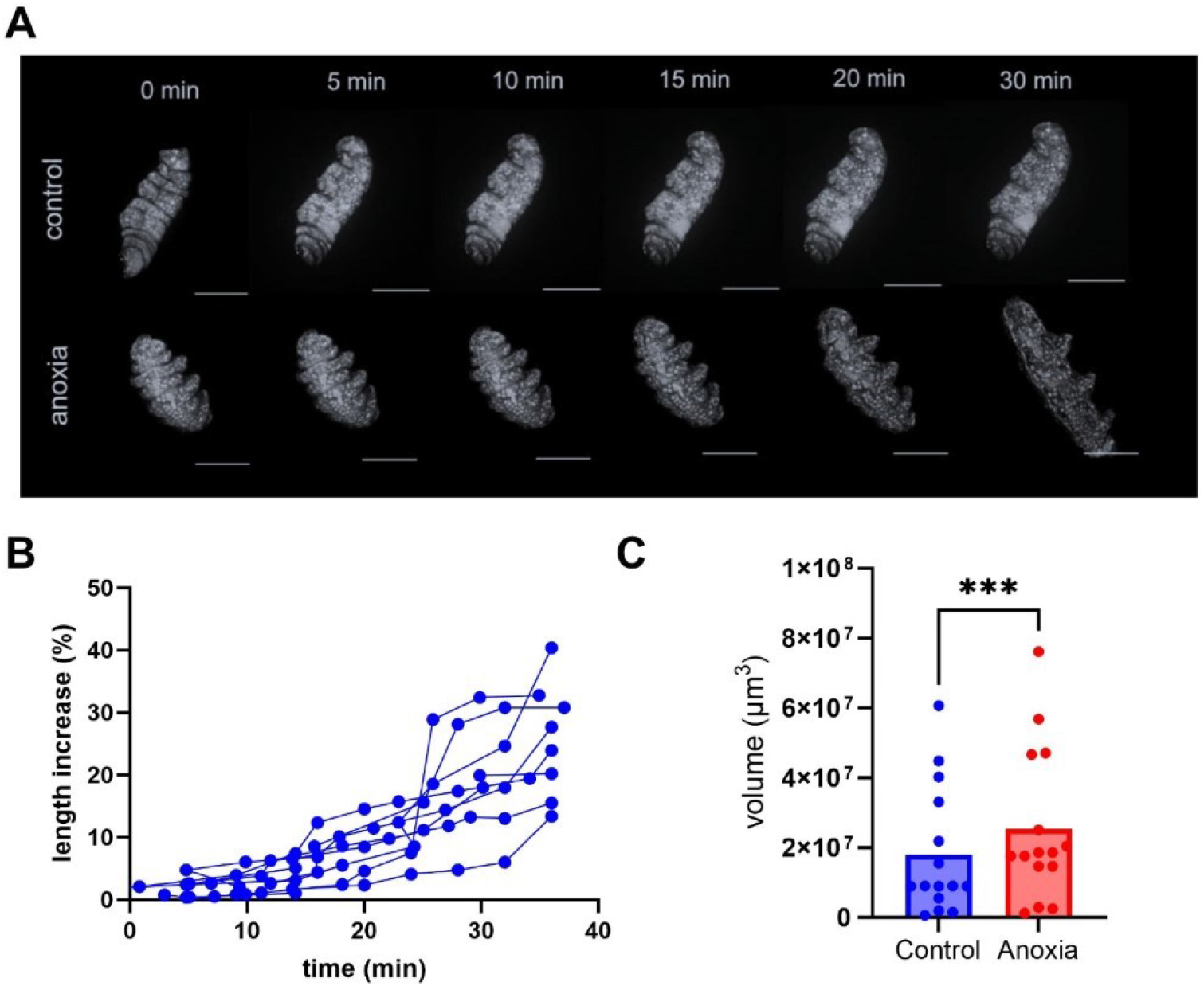
During anoxybiosis the body volume of tardigrade increases. **(A)** Time-lapse imaging and maximum Z-projection visualized the difference between tardigrade in normal oxygen levels and when exposed to anoxia. Scale bar 50 μm. Spinning disk confocal imaging. **(B)** Measurement of body length during anoxybiosis. The length of tardigrade was normalized to its size at the beginning of each movie. n = 8. Used light sheet (FLSM) and spinning disk confocal data. **(C)** Volume measurements of 3D imaged tardigrades. Paired t-test, p = <0.0001, n = 15. FLSM and spinning disk confocal data.

### High-resolution live imaging of tardigrades

Next, we carried-out high-resolution experiments using spinning disk fluorescence microscopy, which provides good resolution with sufficient volumetric imaging performance and imaging speed^24^. The imaging with spinning disk confocal in high-resolution resulted in subcellular level resolution within living tardigrades (Fig. 4). For example, storage cells were clearly visible and characterized by strong NR signal. Storage cells are a specialized cell type of tardigrades, carrying rich lipid energy reserves^25^, which is consistent with observed strong NR signals. Sufficient resolution was required for down-stream image analysis and especially for segmentation and tracking.

**Figure 4.**
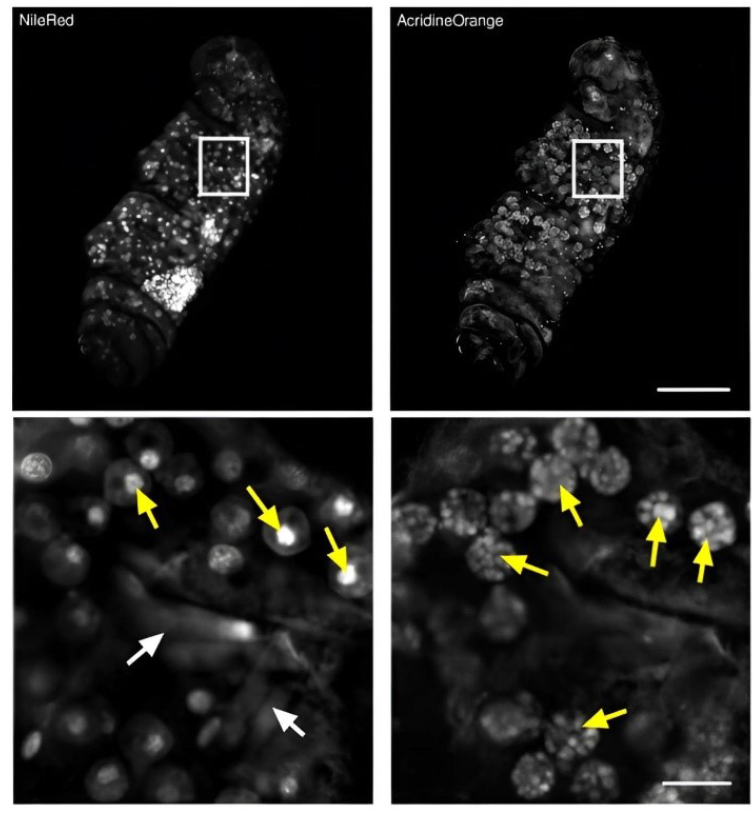
Higher-resolution spinning disk confocal imaging of tardigrades. Spinning disk confocal provided high resolution fluorescence imaging and enabled investigations of change in internal structures of the animal. Yellow arrows annotate storage cells, and White arrows annotate the muscular structures. Scale bar 50 μm / 10 μm.

### Anoxia increases the movement of storage cells

Some studies have showed the importance of storage cells in recovery from environmental stress^26,27^. We hypothesized that storage cells could have an active role during anoxybiosis. To quantitatively analyze cellular movements of storage cells during transition into anoxybiosis, we utilized spinning disk confocal fluorescence time-lapse 3D imaging of tardigrades. From the 3D time-lapse imaging data, we segmented storage cells (stained lipids with Nile Red) and carried out tracking analysis to measure diverse parameters of cellular dynamics of the expanding body of tardigrade (Fig. 5A, supplemental Figure S1). The total number of tracks observed was significantly lower in animals belonging to the control group. This could have resulted from decrease in overall cellular movement in response to oxygen deprivation, segmentation quality or one factor is projected data in order to run the heavy pipelines in Fiji. Besides the amount of tracks we observed that the duration of the tracks was lower in anoxia groups (Fig. 5B). We hypothesized that the passive swelling of the animal would result in increased total distance travelled, increased straight line speed, increased displacement, increased confinement ratio and reduced mean directional change. The total distance travelled (Fig. 5C), mean speed of cells (Fig. 5D) and straight-line speed (Fig. 5E) were increased in anoxia, as expected. However, the measurements of confinement ratio (supplemental figure S2A) or displacement (supplemental figure S2B) did not exhibit statistically significant differences between the anoxia-exposed group and the control group. This suggests that the spatial confinement of cellular movement remained relatively consistent under both conditions. Mean directional change indicates a greater degree of directional changes in cell movement over time reflecting increased cellular exploration or response to environmental cues. Taken together, this data implies that morphological changes in anoxybiosis are not just caused by passive swelling, but may involve active movement of storage cells.

**Figure 5.**
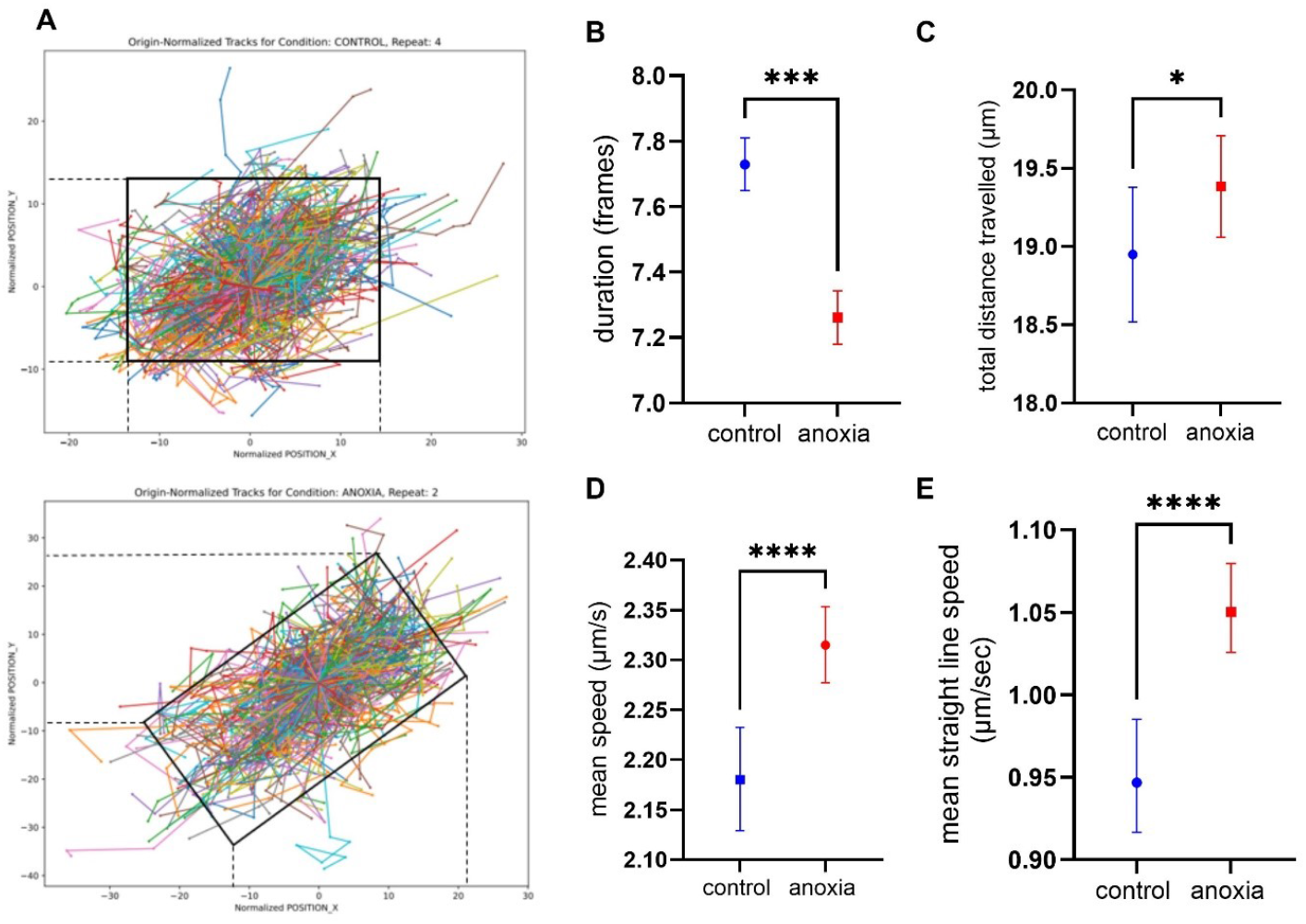
Tracking measurements. All together 5 anoxia and 5 control animals which were exposed to anoxia for 40 minutes, imaged by spinning disk confocal with 4-minute intervals. Tracking was performed with ImageJ for image set that were converted maximum intensity projections to reduce computational burden of the tracking pipeline. Total number of tracks in control group was 1963 and in anoxia the number of tracks was 3657. Data was pooled to reduce the error caused by individuality. **(A)** Visualization of track movement both in control and experimental animal. For the analysis CellTracksColab was used. **(B)** Analysis of duration of tracks. Duration of tracks showed within the anoxia group the tracks lasted shorter period. Mann-Whitney p = 0.0007. **(C)** Analysis of total distance traveled. Longer paths of the cells in anoxia group were observed. Mann-Whitney p = 0.0231 **(D)** Analysis of mean speed. Mean speed (μm/s) was clearly higher in anoxia tracks which was expected. Mann-Whitney p = <0.0001. **(E)** Analysis of mean straight line speed. Mean straight line speed was higher in anoxia Mann-Whitney p = <0.0001.

### Anoxia alters storage cells

To address the hypothesis that storage cells may have active role in anoxybiosis, we analyzed morphology of these cells in detail using high-resolution imaging (Fig. 6A). The analysis of storage cell morphology over time indicated that during anoxia the cell size increased, (Fig. 6B), but their rounded cellular shape did not significantly change (Fig. 6C).

**Figure 6.**
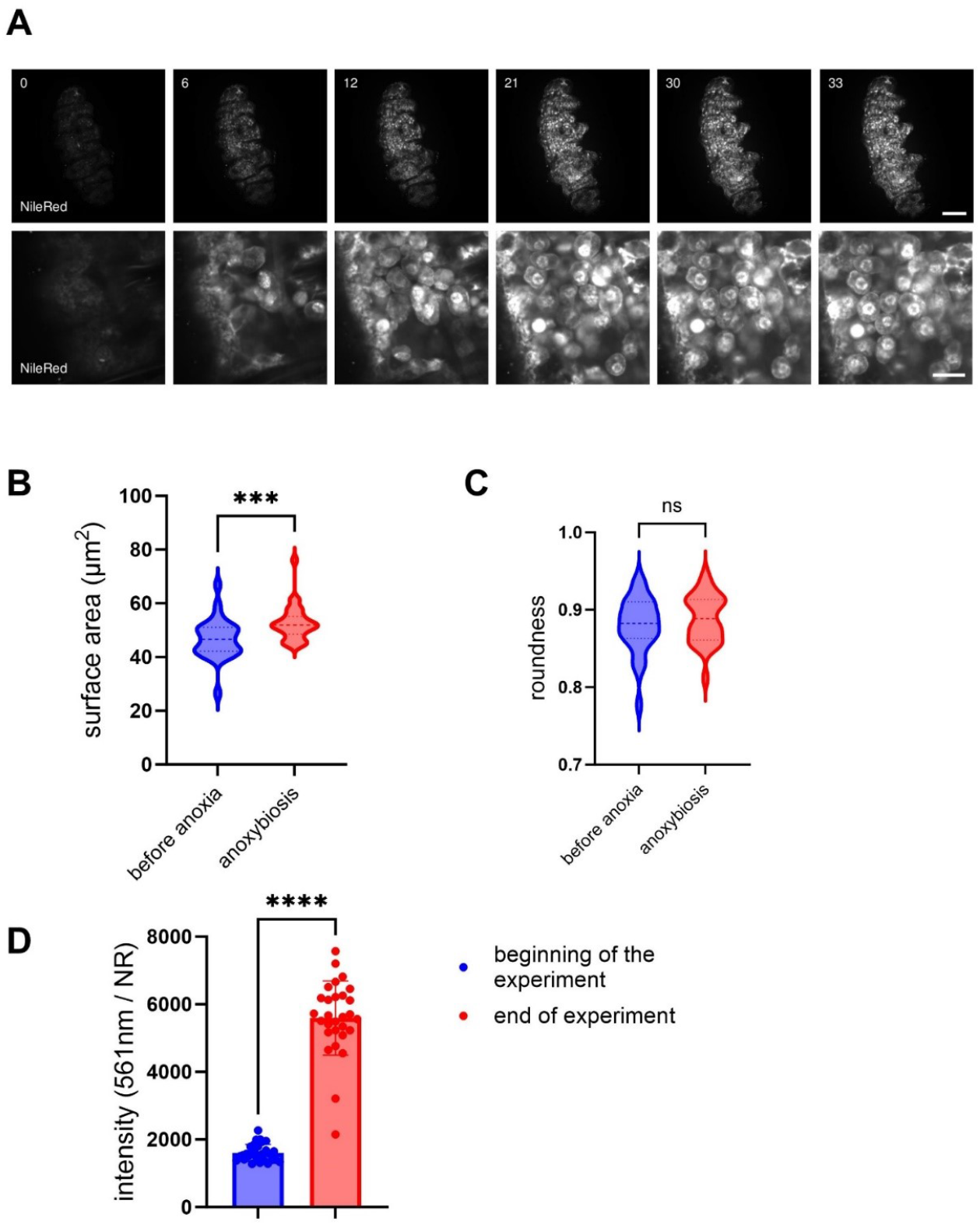
Investigation of storage cells. **(A)** *M. ripperi* stained with Nile Red and Acridine orange imaged with time-lapse high-resolution spinning disk confocal imaging. Whole-animal in upper row and close-up presented in lower row. Scalebar 50 μm in upper row. and 10 μm in lower row. Time in minutes. Five animals were imaged. **(B)** Quantitation of the surface area of storage cells. We measured manually the surface area of the cells, used plane in z-stack which provide the largest cell diameter to be measured, in timepoint zero and in full anoxybiosis. Mann-Whitney test, P = 0.0006. n = 25 (per time point). **(C)** Quantitation of the shape of storage cells. We measured the outline of the cells in timepoint zero and in full anoxybiosis. Roundness was used as quantitative measure for shape. Mann-Whitney, P = 0.45 n = 25 (per time point) **(D)** Measurement of NR fluorescence intensity. Intensity of NR was measured from individual storage cells using excitation at 561nm. Mann-Whitney, P = 0.0001. n = 139 (before anoxia), n = 140 (anoxybiosis).

Serendipitously, we also observed that the NR intensity increased during anoxybiosis (Fig. 6D). NR has interesting solvatochromic properties and it generates stronger red fluorescence when bound to polar rather than non-polar organic solvents^22^. This implied that storage cells underwent metabolic changes during anoxybiosis and their lipid metabolism and constitution was altered. Lipids are typically stored as triacylglycerols (TAG) and metabolized into non-esterified free fatty acids (NEFA) for utilization in beta-oxidation metabolic cycle^28^. This process not only generates energy but also transforms lipids from non-polar form into more polar structures^25,33,^ consistent with increase of NR fluorescence during anoxybiosis. Taken together, our findings illustrated changes of storage cell movement, morphology and composition during anoxybiosis, indicating active role of storage cells during anoxybiosis.

## Discussion

In this study, we investigated the morphological changes of tardigrades during anoxybiosis by developing live fluorescence imaging protocols and using high-resolution and intravital imaging technologies.

First, we observed that two tardigrade species *P. fairbanksi* and *M. ripperi* had different sensitivity to anoxia. This is in-line with previous studies have found that anoxybiosis in tardigrades is species specific^14,29^. For example, the *Richtersius cf. coronifer* and *Hypsibius exemplaris* responded to hypoxia differently: Both species exhibited irregular body movements at similar oxygen levels, but larger fraction of *R.cf.coronifer* regained regular movements after 10 hours of recovery from hypoxia ^6^.

The utilization of high-resolution imaging has afforded us an unprecedented level of detail in observing the structural alterations in tardigrades undergoing anoxybiosis. These techniques have demonstrated the preservation of cellular integrity and the morphological adaptations that can explain tolerance for anoxic conditions. The observed changes in cuticle structure (expansion, swelling), the reorganization of cells (storage cell movement), and the alterations of metabolism within storage cells (lipids) may be linked to survival mechanisms. Studies that have investigated cryptobiosis in general have pointed out storage cells and their function related to stress tolerance^29,30^. The storage cells are considered to function as the energy storages of tardigrades^25,27,31^, but these cells may have also other roles such as in vitellogenesis^32^. In anhydrobiosis, there is increased expression of genes belonging to the SAHS (secretory abundant heat-soluble) family in storage cells hinting at their potential involvement in desiccation tolerance^21^. Our experiments implied changes in tardigrade metabolism in anoxybiosis, indicated by intensity changes of NR dye. Advanced imaging modalities such as mass spectrometry imaging or CARS (Coherent Anti-Stokes Raman Scattering) microscopy could later provide deeper information of metabolic responses of tardigrades to anoxic conditions. The importance of further investigation of tardigrade metabolism has been emphasized^6^ as it still has many open questions.

We developed protocols for live imaging of tardigrades using fluorescent dyes and high-resolution imaging. On the other hand, recent work has described protocol to make fluorescent transgenic tardigrade models^21^, which can facilitate future studies of molecular functions and interactions during cryptobiotic processes. During the preparation of this manuscript also the use of some other fluorescent dyes in imaging of tardigrades were reported^33^. In addition, fluorescence shadow imaging can be utilized to image external morphology of tardigrades^34^. These methods are complimentary to each other, and together expand the experimental toolbox available to study tardigrade biology^35,36^. By correlating detailed structural observations with dynamic physiological processes, we can begin to construct a comprehensive picture of how tardigrades manage to halt metabolic processes and protect cellular components during anoxia, only to resume normal physiological functions upon reoxygenation.

Intravital imaging complemented high-resolution static imaging by providing a dynamic view of the tardigrade’s response to anoxia. Techniques such as live-cell fluorescence microscopy have enabled the real-time tracking of physiological processes, including the redistribution of intracellular components^37^. Moreover, intravital imaging allowed for the study of tardigrade behavior and physiology in a more natural context, facilitating future studies on how environmental factors influence the onset and termination of anoxybiosis. The insights gained from our study extend beyond the biology of tardigrades, offering valuable lessons for the broader field of extremophile research. Elucidation of survival mechanisms in inhospitable conditions is relevant not only for evolutionary biology and ecology but also has potential applications in biotechnology and medicine^31^. It has been proposed that the understanding mechanisms of stress tolerance could lead to innovative strategies for preserving cells and tissues in medical settings or developing life-support systems^38^.

In conclusion, the utilization of high-resolution and intravital imaging in the investigation of tardigrade anoxybiosis facilitated the acquisition of novel insights that may elucidate the underlying mechanisms of extreme survival strategies. Our findings also support further study of the components that take part when tardigrades are exposed to different oxygen levels. Because anoxybiosis differs from cryptobiotic forms that rely on tun formation, we consider the deeper understanding of what happens with storage cells during and after anoxybiosis vital. As these imaging technologies are harnessed and further developed, it enables uncovering new biological wonders and deepening our understanding of the resilience of life in the face of adverse conditions.

## Materials and methods

### Tardigrade culture

*Macrobiotus ripperi* collected from moss in Jyväskylä, Finland in 2014. Animals were cultured in high wall plastic petridish (Ø50mm, height 20.3mm) (Sterilin™ Petri Dishes, Thermo Scientific). The dish was scratched with sandpaper for the animals to be able to walk on the surface. Spring water (Kotimaista lähdevesi, Multiala spring, Finland) was used as medium, and animals were fed with RG Complete (Planktovie SAS, Marseille, France) and young *Panagrellus pycnus* nematodes (laboratory cultured, batch gained from University of Jyväskylä). Incubator was set on 18°C. No light-dark cycle was applied. Animals were imaged weekly with Zeiss Stemi 350 stereomicroscope (Carl Zeiss AG, Oberkochen, Germany) to monitor culture quality, feeding and cleaning.

### Testing the anesthetic

Testing the anesthetics was performed by applying the anesthetic on agarose mounted tardigrades. The effect was observed and imaged by Zeiss AxioZoom stereomicroscope. All together 25 *M. ripperi* and 23 *P. fairbanksi* were used at timepoints: 15 minutes, 30 minutes, 1 hour and 4 hours. Concentrations of anesthetic solutions were based on literature values with Levamisole and MS222 concentration was same as used with fish embryos.

### Tardigrade staining, anesthesia and anoxia solution

The animals were stained overnight in a total volume of 2ml of staining solution. The dyes were prepared by diluting 5 μM acridine orange (ThermoFischer Scientific, Germany) in spring water and 10 μM NileRed (Sigma Aldrich, Germany) in DMSO. Prior to imaging, the animals were rinsed in order to eliminate any residual dye. Acridine orange was employed for comprehensive labelling of the nuclei. Additionally, the dye imparts a yellowish hue to the entire animal, which facilitates detection, particularly in light sheet imaging. Nile Red was used to stain lipids in storage cells. The dyes exhibit slight spectral overlap, yet this did not necessitate any adjustments in our studies due to their structural specificity.

The animals were mounted in low-melting-point agarose (Sigma Aldrich, Germany) and positioned on either a silicon sample holder for light sheet imaging or a glass bottom dish with inverted microscopes, such as a spinning disk confocal. The animals were anaesthetized with 2 g/l tricaine (MS222, Sigma Aldrich, Germany). This was selected over levamisole due to the higher survival rate. Once the mounting process was complete, the animals were ready for imaging. Following imaging, the animals were transferred to fresh spring water to recover.

For chemical anoxia, sodium metabisulfite (JT Baker, ThermoScientific, Germany) was used. The concentration of sodium metabisulfite in the anoxia solution was 1mg/ml. Oxygen levels of the solution were tested with JBL ProAquatest O2 (JBL GmbH & Co. KG, Neuhofen, Germany), and was at/below detection limit (<0.2mg/ml) of the test. Sodium metabisulfite decreases the pH which corrected to match the spring water (pH 6.5) with 1M NaOH. Anoxia solution was pipetted on the agarose in large excess.

Control groups were handled the very same protocol except the chemically applied anoxia exposure.

### Light-sheet imaging

Light sheet fluorescent microscope (M2 Aurora Airy beam, M Squared Life, Glasgow, UK) was used in intravital imaging of tardigrade anoxybiosis. Tardigrade was mounted on a single drop of agarose on a silicon sample holder and set in the imaging chamber filled with anoxia solution (J.T.Baker, FisherScientific, Germany). Tricaine was used to anesthetize the animal before anoxic treatment to make sure the position is beneficial for the imaging.

Excitation and detection objectives (Special Optics) provided magnification and numerical aperture value based on imaging media. The media was water / agarose. The refractive Index (RI) of water was 1.33, reaching magnification up to 16x with NA0.4 and resolution up to 2.35 pixels / μm, optical resolution of 0.6-0.7 μm, with up to 870μm x 870μm full field-of-view. In some cases, we used smaller field-of-view for optimal data collection. In time lapse imaging, a full 3D-stack was collected with 4-minute intervals for 40 minutes.

### Spinning disk imaging

Spinning disk (multi-point) confocal microscope (3i Marianas CSU-W1 spinning disk, Denver, Colorado, US) was used to gain high resolution images of tardigrades. Sample preparation included agarose mounting on a glass bottom dish, anesthesia and anoxia treatment with experimental animals. Imaging settings were straight forward; excitation lasers 488 nm and 561 nm, fixed filters and camera Hamamatsu sCMOS Orca Flash4.0 2048 × 2048 pixels, 6.5×6.5μm and 40x NA1.1 Zeiss LD C-Apochromat WI objective with working distance of 620 μm. Optical resolution with 561nm laser reaches up to 0.225 μm. With 40x objective our FOV is 325μm x 325 μm so one animal fit in, camera resolution up to 6.3 pixels/μm. Time lapse imaging was set to 4-minute intervals for 40 minutes.

### Stereomicroscope imaging assays

Motility studies carried out with Zeiss AxioZoom.V16 stereomicroscope with 1×0.125 objective (Carl Zeiss AG, Oberkochen, Germany). Short bright field videos were captured with Zeiss movie maker and tardigrades manually tracked using Manual Tracking (Fiji /ImageJ, Wayne Rasband and contributors, NIH, USA) in the presence and absence of anesthetic. Initial testing of fluorescent dyes was done with Zeiss AxioZoom using filters Alexa 488 - filter set 38 HE and Alexa 568 - filter set 45.

### Cell tracking

For cell tracking of spinning disk data, we used Fiji Trackmate^39^. With larger data sets (i.e., light sheet) we used Zeiss Arivis Vision 4D 3.5 (Zeiss Group, Oberkochen, Germany) for particle detection and tracking. CellTrackColab^40^ was used for deeper analysis of the Trackmate defined tracks of confocal spinning disk images sets.

In Fiji Trackmate, we used LoG detection for 2×2 binned image sets. Estimated object diameter was set 10 μm and threshold 30; these values were used for every image sets. For tracking Simple LAP tracker, max distance was set to 30 μm, same as gap closing. Max frame gap was set to 2, however the 2-spot tracks were filtered out in later analysis steps.

In Arivis we used analysis panel to create pipeline for tracking cells and particles. Common workflow included setting up the detection for desired channels (lipids for Nile red and overall tracking with AO). Additional parameters included background correction, morphology filter and shape detection. To improve segmentation, we also applied either intensity-based watershed or machine learning based segmentation.

### 3D volume measurements

3D renderings for volume measurements were done using Imaris (Bitplane, South Windsor, CT, USA). After image file format conversion, the Surface module was used and threshold was set manually. 3D renderings were carried out with Zeiss Arivis as well.

### Statistical analyses

The statistical analyses were conducted using GraphPad Prism (version 10.1.2 for Windows, GraphPad Software, Boston, Massachusetts, USA; www.graphpad.com). Non-parametric Mann-Whitney analyses and t-tests were conducted for non-normal and normally distributed data, respectively. The survival data were analyzed using the Mantel-Cox log-rank test. The CellTrack Colab data were analyzed on the server and randomizations were based on the experimental nature of the data. The image files were packed in max projection format for lighter processing. The settings were consistent across all data sets, ensuring that any errors were adjusted correctly for the entire data set.

## Acknowledgements

We would like to express our gratitude to Cell Imaging Core and Zebrafish Core of Turku Bioscience Centre, supported by Biocenter Finland, for their invaluable support, infrastructure and resources.

## Competing interests

University of Turku has registered trademark 3DFLUOHISTO®, and I.P. is involved in the commercialization of 3DFLUOHISTO-technology. Other authors declare that they have no conflict of interest.

## Author contributions

Conceptualization: AMHS, SC, IP

Methodology: AMHS, IP

Validation: AMHS

Formal analysis: AMHS, IP

Investigation: AMHS

Resources: SC, IP

Data Curation:

Writing – Original Draft: AMHS, IP

Writing – Review & Editing AMHS, SC, IP

Visualization: AMHS

Supervision: SC, IP

Project administration: IP

Funding acquisition: SC, IP

## Funding

University of Turku (I.P.), BusinessFinland (I.P.), Finnish Foundation for Cardiovascular Research (I.P.),

## Data availability

Data is available from authors upon reasonable request.

