## Supplemental Figure S1 for "High resolution live imaging of tardigrade response to anoxia"

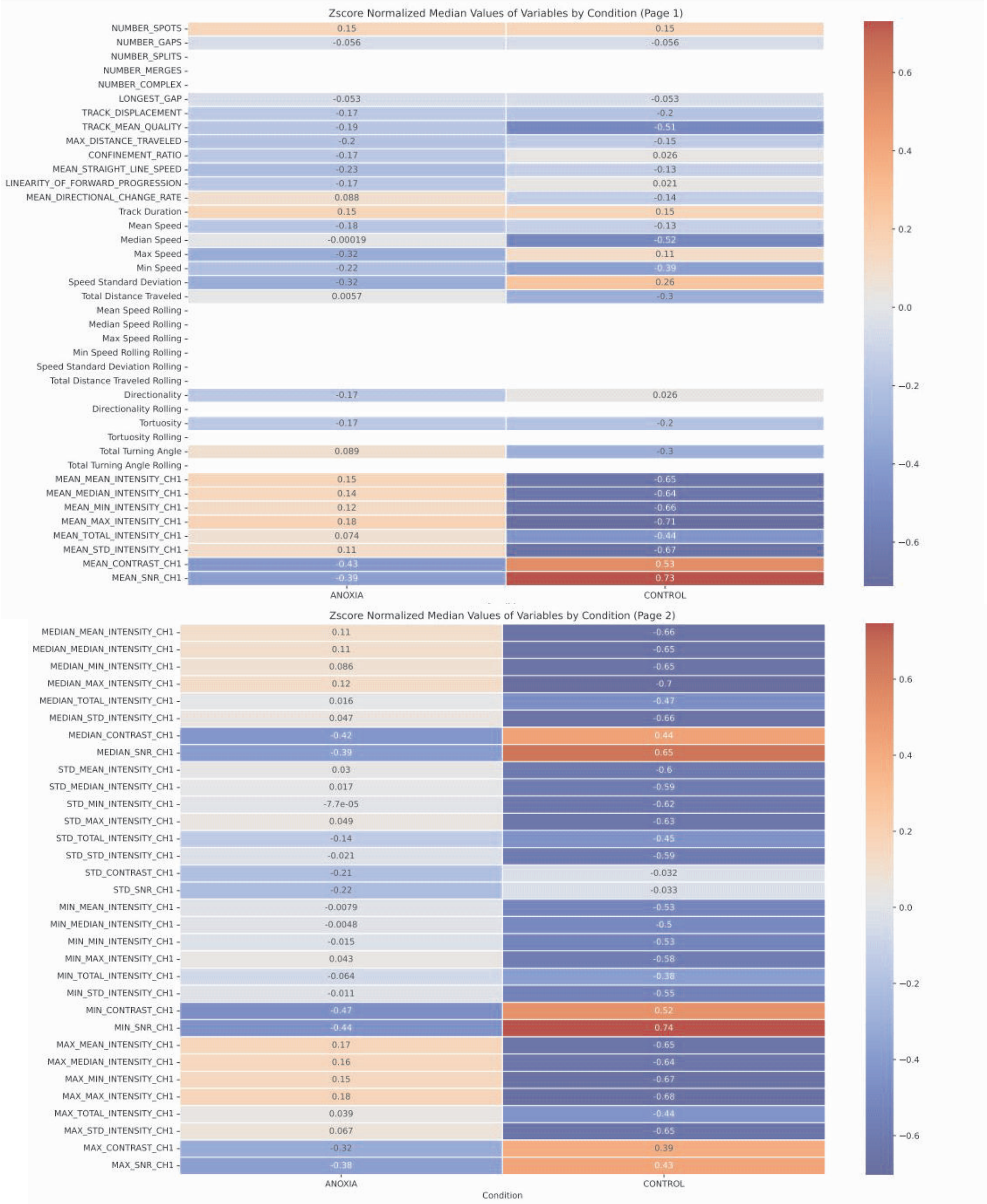


**Figure S1.** Tracking parameters between the control and anoxia esposed groups. Randomized and normalized data was sorted as a heat map to illustrate the difference between groups. This was created by CellTrackColab and tracking information was run via Trackmate.
