## Supplemental Figure S2 for "High resolution live imaging of tardigrade response to anoxia"

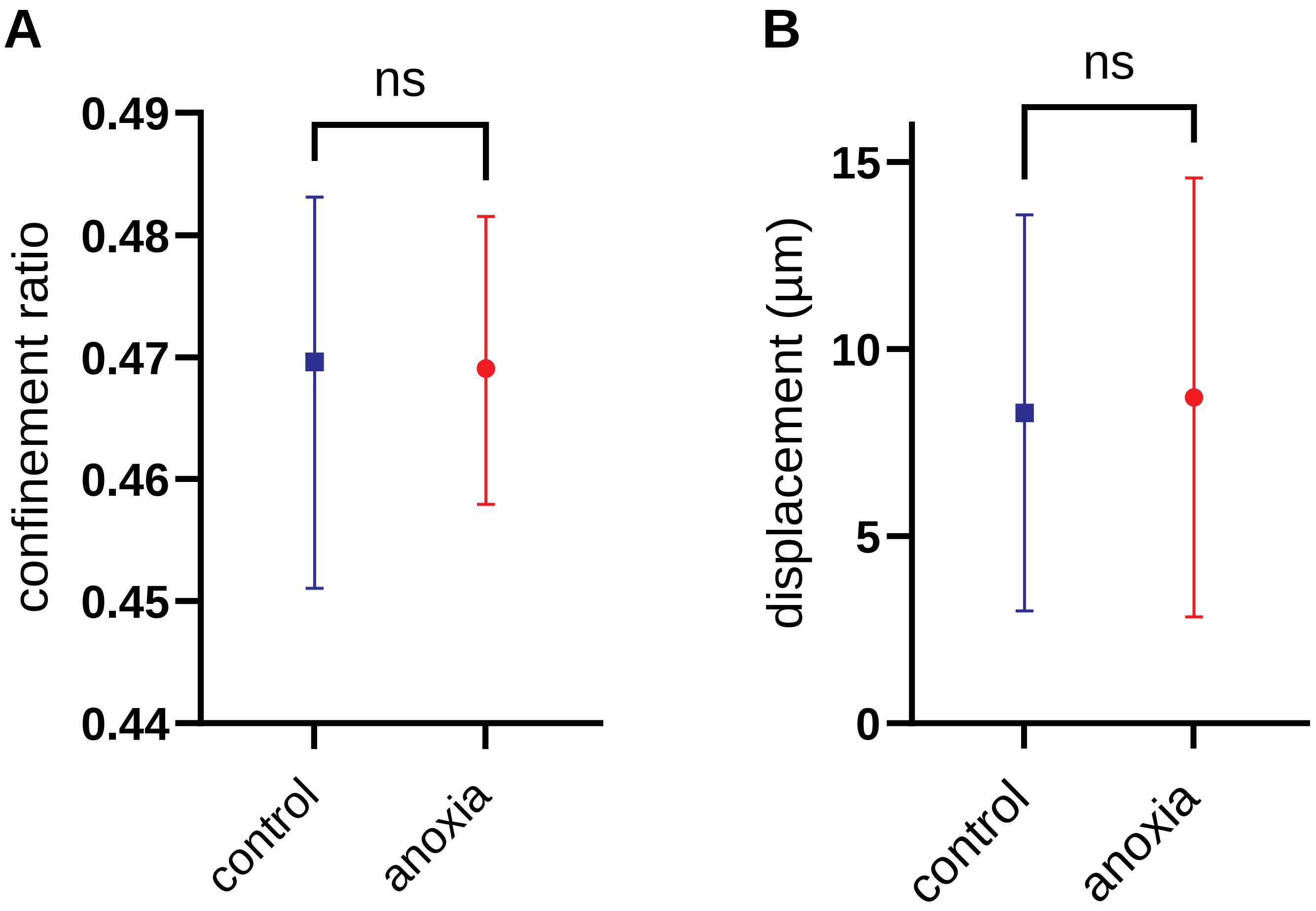


**Figure S2.** Major tracking parameters. (A) Confinement ratio shows the how still the movement is. The closer to 0, the less the particles has moved away from its starting point. There was no statistical significange between control and experimental group (p = 0.2407, Mann-Whithey), however the value indicates that in both cases the cells have travelled during the period of time. (B) Displacement illustrates distance between starting and end point (µm), however it does not take account the total distance the particles has travelled. No statistical signifigance between the groups (p = 0.0887, Mann-Whitney).
